# Colour Preferences of UK Garden Birds at Supplementary Seed Feeders

**DOI:** 10.1101/103671

**Authors:** Luke Rothery, Graham W. Scott, Lesley J. Morrell

**Affiliations:** School of Environmental Sciences, University of Hull, Kingston-upon-Hull, UK

## Abstract

Supplementary feeding of garden birds has benefits for both bird populations and human wellbeing. Birds have excellent colour vision, and show preferences for food items of particular colours, but research into colour preferences associated with artificial feeders is limited to hummingbirds. Here, we investigated the colour preferences of common UK garden birds foraging at seed-dispensing artificial feeders containing identical food. We presented birds simultaneously with an array of eight differently coloured feeders, and recorded the number of visits made to each colour over 370 30-minute observation periods in the winter of 2014/15. In addition, we surveyed visitors to a garden centre and science festival to determine the colour preferences of likely purchasers of seed feeders. Our results suggest that silver and green feeders were visited by higher numbers of individuals of several common garden bird species, while red and yellow feeders received fewer visits. In contrast, people preferred red, yellow, blue and green feeders.

## Introduction

It has been estimated that 20-30% of people in more developed areas of the world provide wild birds with additional food (supplementary feeding) at some point in the year (typically during the winter months) [1,2]. In the UK, approximately 60% of households with gardens provide food for birds [3], estimated at 12.6 million households [1], 7.4 million of which use bird feeders [4]. As a result the UK wild bird feeding industry was estimated as being worth £210m per annum [5], and the wild bird care market rose 15% in value between 2014 and 2015 [6]. Levels of bird feeding vary enormously across society [7], but the importance of the connection between people and nature to human well-being in urban environments is well established [8]. People feed birds because it gives them a sense of personal wellbeing, although the underpinning emotions, experiences and personal perceptions of the people feeding birds are certainly more complex than such a simplistic statement might suggest [9]. Some people (those involved in avian monitoring or research) feed birds in order to attract them for capture, measurement and subsequent release.

During the northern hemisphere winter natural food resources are at their lowest level of availability [10] and a bird’s thermodynamic costs are at their highest [11]. Over winter survival is thus highly dependent upon the characteristics and availability of food supply [10]. Gaining enough energy each day to ensure overnight survival is particularly important for small passerines: individuals in the tit family (Paridae) can lose up to 10% of their body weight overnight in winter [12]. Supplementary feeding may off-set the effects of winter resource depletion [13] and in many cases a winter feeding station may be the most abundant and dependable food source in a particular area [14]. Supplementary feeding has been recorded as having a number of other benefits to birds, including larger clutch sizes (house sparrows *Passer domesticus* [15]), better body condition and more rapid recovery from injury (Carolina chickadee *Parus carolinensis*, tufted titmice *Parus bicolor* and white-breasted nuthatch *Sitta carolinensis* [16]). Supplementary feeding increases both the range of species and number of individuals visiting gardens [1,17] and increases abundance at a landscape scale [1]. In the UK, for example, supplementary feeding has been implicated in population increases of both house sparrow and starling (*Sturnus vulgaris* [18]) and may be important in the evolution of ‘new’ migration strategies amongst over-wintering blackcap (*Sylvia atricapilla* [19]).

In order that the benefits of supplementary feeding to both birds and the people who feed them are realised it is essential that food be provided in a way that makes it accessible to birds. In the case of the seed based foods provided to passerines supplementary feeding most often involves the use of commercially available tubular seed dispensers. These feeders commonly consist of a transparent plastic tube through which seeds are visible to birds and coloured metal or plastic lids, bases, perches and feeder ports. Here, we report an investigation into whether the colour of these metal or plastic parts affected the number of birds choosing to feed at a particular feeder. For a feeder to attract larger numbers of birds, something likely to be seen as preferable by those that purchase feeders, the colour should be attractive or neutral to either a particular target species, or seed feeding birds more generally. A feeder the colour of which birds avoid would not be an effective feeder.

Birds have excellent colour vision and exhibit the ability to distinguish and choose between different colours and shades [e.g. 20–22]. Here, we focus on colour preferences in relation to foraging. Multiple studies report preferences of birds for food items of a particular colour. Great tits (*Parus major*) blue tits (*Cyanistes caeruleus*) and Eurasian nuthatches (*Sitta europaea*) all preferred uncoloured (natural) peanuts over those that had been dyed white [23]. Willson et al. [24] reviewed a number of studies demonstrating that frugivorous birds prefer black or red grapes or cherries over other colours such as green and yellow, but point out that preference for colour here is confounded by preference for other factors associated with colour, such as ripeness, size and nutritional value [24]. Other studies have used artificial or novel foods dyed different colours and found colour-based preferences [24–26]. Willson et al [24] reported a preference for red, and avoidance of yellow in three species of frugivorous bird, while North Island robins (*Petroica longipes*) preferred yellow and avoid blue and brown [26] for example.

Preferences for colour associated with supplementary feeders, rather than food, have exclusively focused on the preferences of hummingbirds (Trochillidae) at feeders designed to provide sugar syrup. While hummingbird-pollinated flowers tend to be red [27,28], and birds tend to prefer red-pigmented flowers over those lacking red pigments (e.g. [29–31], reviewed in [28]), experimental studies on feeders do not show a consistent preference for any particular colour (e.g. [32–34], reviewed in [28]). Instead, factors such as location [35,36], previous experience [37–39] and nectar quality [35,39] appear to be more important in determining choice. We have been unable to find any peer-reviewed studies of the impact of seed dispensing feeder colour on bird feeding behaviour. One anecdotal report [40] suggested that work carried out by the British Trust for Ornithology demonstrated colour-based preferences for birds visiting seed and peanut feeders, namely that blue seed feeders are preferred during the summer, while silver feeders are preferred in winter (although goldfinches preferred green), and red peanut feeders are preferred over other colours. The primary aim of our research was to investigate the effect of feeder colour on the feeding preferences of wild birds. As an additional aim we investigated the level to which birds and the humans who feed them agreed on their preferred feeder colour, an important consideration for those who make and sell feeders and those who use them.

## Methods

### Bird colour preference

To explore the effect of colour on the number of visits by birds, we recorded bird visit rates to 8 different coloured feeders at three sites on 78 sampling days during the winter/spring of 2014/15 (November 2014 to May 2015).

Data were collected at Tophill Low Nature Reserve (Driffield, East Yorkshire TA 075,492), The University of Hull Botanic Garden (Cottingham, East Yorkshire TA 050,329) and a suburban garden in Otley (West Yorkshire SE 195,472). These sites were chosen due to accessibility and the presence of existing artificial feeders with regular avian visitors. The feeders used (Natures Feast Royal Seed Feeders, Westland Horticulture) were of transparent tubular design with metal lids, two metal ports and two straight metal perches. The metal parts of each feeder were painted a single colour using Hammerite Metal Paint. The proprietary colours used were *Smooth Black, Smooth Blue, Smooth Dark Green, Smooth Red, Smooth White, Smooth Yellow, Hammered Silver* and Purple (achieved by mixing *Smooth Blue, Smooth Red* and *Smooth White* at a ratio of 3:2:1). Analysis of the feeder colours can be found in the section below. Throughout the experiment the feeders were filled with “Nature’s Feast High energy No Mess 12 Seed Blend” (Westland Horticulture, UK).

At each site the feeders were suspended in a line from a metal cross-bar, 30 cm apart from one another and 1.5m above the ground. At any time, one feeder of each of the 8 colours used was available (see supporting information: Fig S1). The order of the feeders along the cross-bar was changed after every 30 minute observation period according to a pre-determined random pattern to control for any preferences based on feeder position rather than colour. Feeders were filled at the beginning of each observation period and cleaned thoroughly every 14 days. During each data collection session the numbers of feeding visits by birds to each of the feeders in the array were recorded over 30 minutes. A feeding visit was defined as a bird landing on the perch and taking food from the feeder port. Birds were identified to species level, but as it was not possible to distinguish between individuals of the same species, each visit to the feeders was counted as an independent data point. All observations periods were video recorded (Sony Handycam HDR-CX240E) mounted on a tripod approximately 10m from the feeders. Identification and counting of birds either took place in real-time in the field or later using the video recordings (where the number of visits was too high to allow for accurate real-time recording).

Data were collected across a total of 370 observation periods (Otley: 208; Tophill; 26 Botanic Gardens: 136), and a total of 7535 visits to the feeders were recorded (table 1).

**Table 1:**
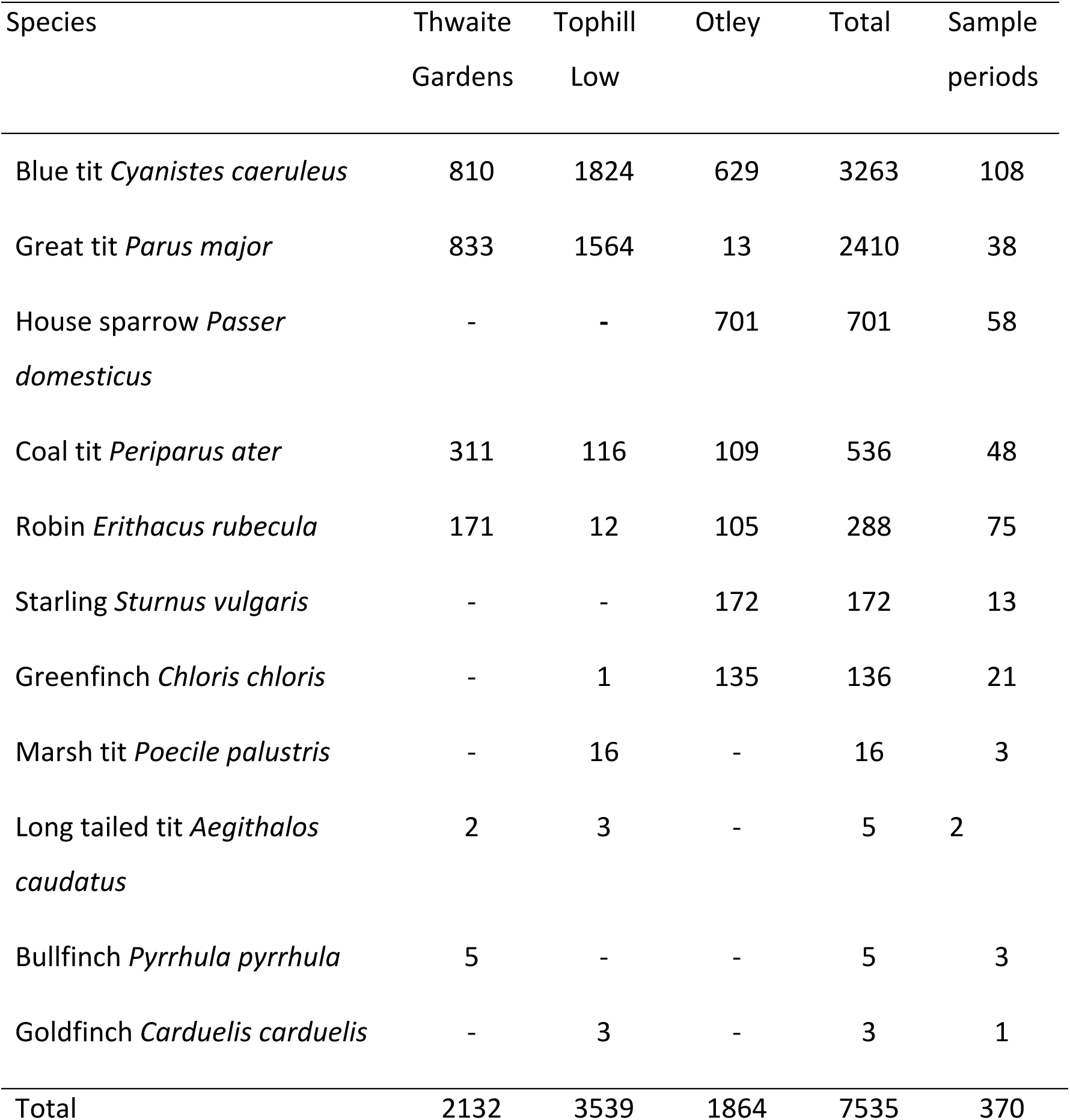
Summary of data, showing the total number of visits by each species at each site, and the number of sample periods in which that species was observed at least once.

### Human colour preference

To assess the preferences of likely purchasers of bird feeders, we collected data in a garden centre (Hornsea Garden Centre, Sigglesthorne, Hornsea, UK) where similar feeders were sold (3 days, 8 2-hour sample periods) and at the University of Hull Science Festival (1 day as a single sample period). At each venue we explained to adult volunteers that we were investigating the choices made by birds and people but we did not provide any information on actual bird preferences (supporting information: Fig S2). People were shown the coloured feeders used in the study and asked simply to indicate (by placing a token in an appropriately coloured container) which they would be most likely to buy for their own garden. Containers were emptied and tokens counted at the end of each sample period. In total, 587 ‘votes’ were cast during the poll.

### Data analysis

All analysis was carried out using R v3.2.3 [41]. The total number of visits (across all species, to give a measure of the overall preference for particular colours) to the feeders were analysed using a generalised linear mixed effects model with a Poisson error distribution (as appropriate for count data). Observation period and site were added as random effects to account for non-independence of visits to feeders displayed at the same time, and overall differences in bird populations at a given site. An observation-level random effect was included to account for overdispersion in the data [42]. Re-leveling the data within the model allowed for all pairwise comparisons between colours to be made, and p-values were corrected for multiple testing across pairwise comparisons using the false discovery rate control method [43]. The same analysis was used for the number of visits for each species with more than 100 total visits to the feeders (see supplementary tables S1-S5), to evaluate whether different species had different colour preferences. Preferences expressed by visitors to the garden centre and science festival were also analysed using the same methodology.

### Feeder colour analysis

To objectively describe the colour of the feeders, photographs of the feeder lids were taken in RAW format using a Canon Powershot G12 camera. Lids were placed into a light tent (EZCube, Ventura, CA, USA) under daylight spectrum illumination with a white reflectance standard (Ocean Optics, Dunedin, FL, USA).

Images were processed using the Image Calibration and Analysis Toolbox [44] plugin for ImageJ 1.50i [45]. After using the toolbox to linearise and standardise the image against the white standard, a patch on each feeder that was approximately the same distance and orientation as the reflectance standard and free from specular reflections, was selected, and the mean camera-specific RGB values of the patch were recorded (16-bit colour depth).

To summarise the luminance independent colour measures, RG and BY ratios were calculated (RG = (R-G)/(R+G); BY = B–((R+G)/2)/ B+((R+G)/2); Fig 1A). These ratios describe the redness versus greenness (RG), and blueness versus yellowness (BY) of a stimulus and approximate human and potential avian opponent colour channels [46]. Additionally, luminance ((R+G+B)/3) is shown in Fig 1B. As the camera was not UV sensitive and had not been characterised (i.e. the spectral sensitivity of each sensor measured), it was not possible to measure reflectance in the UV range or transform the RGB values into avian colour space [44].

**Fig 1:**
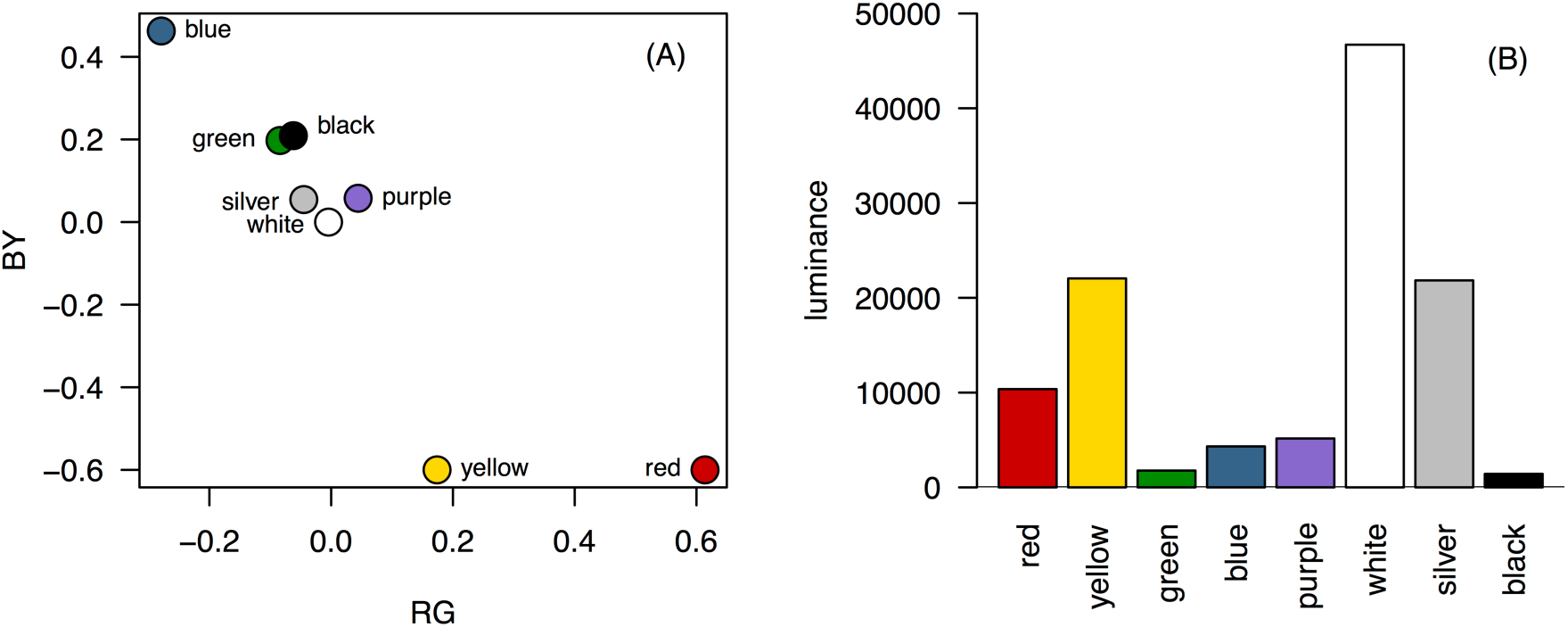
Analysis of feeder colour. (A) RG and BY ratios, and (B) luminance for the 8 different feeder colours

### Ethical statement

Experiments were approved by the University of Hull’s School of Biological, Biomedical and Environmental Sciences and Faculty of Science and Engineering ethical review committees before commencement. All avian work was observational, and carried out at locations where supplementary feeding of birds already occurred and would continue after data collection was completed. Permission to carry out fieldwork was granted by the University of Hull (Thwaite Gardens), Richard Hampshire (Tophill Low Reserve Warden) and Mark Rothery (Otley site owner). The field studies did not involve endangered or protected species. All participation in the human colour preference was entirely voluntary and the purpose of the experiment was explained to the participants either verbally or via an A4 poster displayed near the stand (Fig S2). Written consent was not obtained to ensure participation was simple and to maximise the number of participants, and approved by the institutional review boards above. No data on the participants (other than their choice of colour) was collected.

## Results

### Bird colour preference

There was a significant effect of feeder colour on the number of visits to the feeders (F_7,875_=6.120, p < 0.001; Fig 2A). Birds made significantly more visits to the silver feeder and significantly fewer visits to the red and yellow feeders than any other colour (all p<0.05; table 2). Green was visited significantly more often than any other colour except silver, but there was no difference in the number of visits to blue, purple, white and black.

**Table 2:**
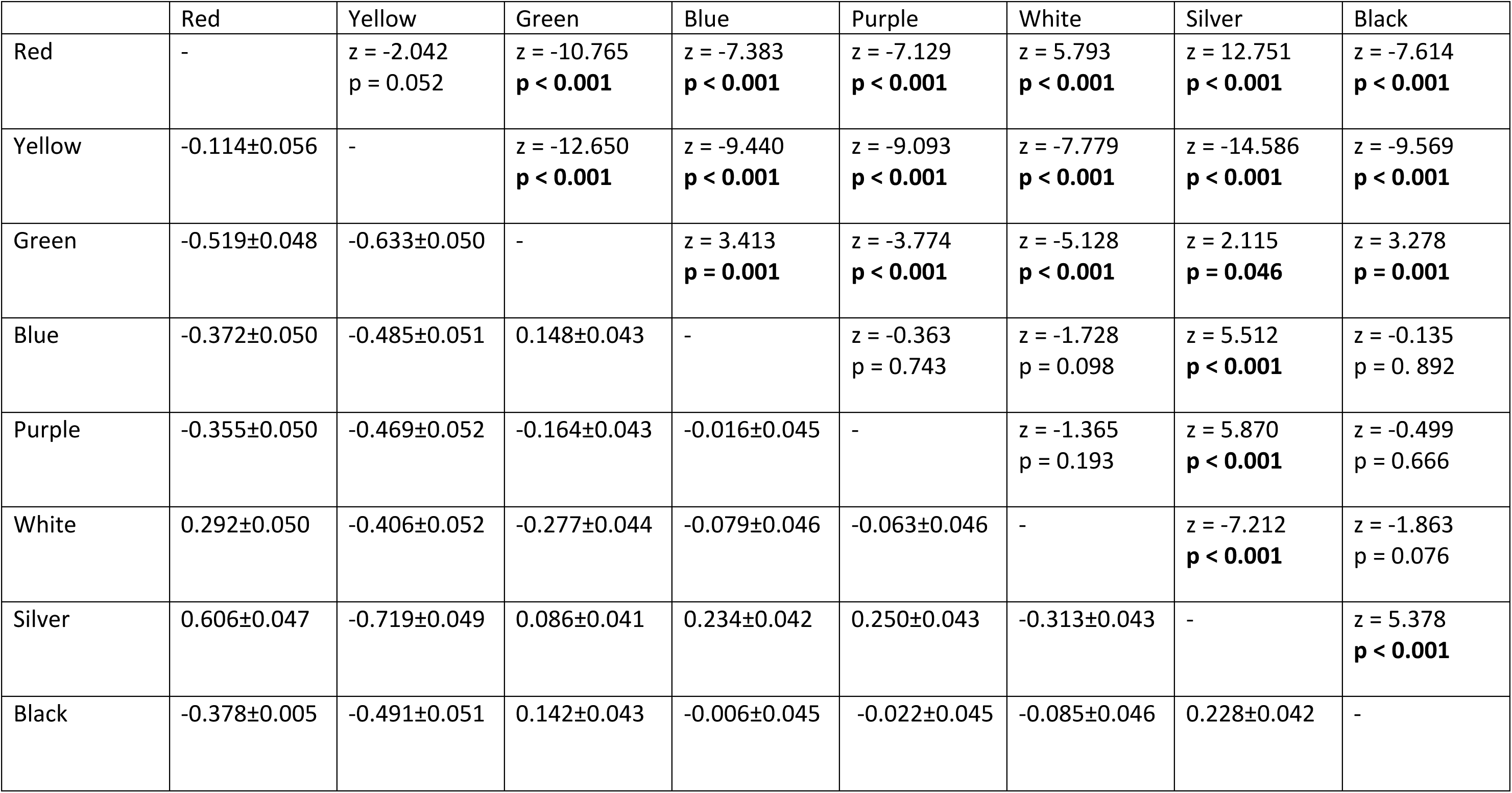
Pairwise comparisons of visits to feeders.

The cells above the diagonal show the z- and p-values, while the estimate ± standard error is below the diagonal. Significant p-values are highlighted in bold. Adjusted p-values following false discovery rate control are presented.

For blue tits (Fig 2B, supporting table S1), there was a significant effect of colour on number of visits (F_7,749.73_=4.3575, P < 0.001). Yellow and red were the least visited colours, and were visited with similar regularity (table S1, p > 0.05). Yellow was visited significantly less than all other colours except white (p > 0.050), while visits to red were not different from white or green (p > 0.05). There were no differences in the number of visits between the other colours (p > 0.05; table S1).

For great tits (Fig 2C, table S2), there was a significant effect of colour on number of visits (F_7,259_ = 2.671, p = 0.011). There were significantly fewer visits to yellow than to all other colours except red (p < 0.05 in all cases), while red was visited significantly less often than green (p = 0.017). There were no other significant pairwise differences (table S2).

For coal tits, there was a significant overall effect of colour on visits (F_7,329_=3.796, p < 0.001), but no significant pairwise comparisons were found after correction for multiple testing (Fig 2D; table S3).

For house sparrows, there was a significant effect of colour on visits (F_7,399_ = 11.139, P < 0.001). The yellow feeder was visited significantly less often than all other colours (Fig 2E, table S4, p < 0.05 in all cases), and red was visited less often than blue, green, silver and black (p < 0.05). White and purple were visited less often than blue, green and black (p < 0.05) which were the colours visited most (although not significantly more than silver; table S4)

For robins, there was a significant effect of colour on visits (F_7,518_=3.1033, p = 0.003; Fig 2F). Black, the most visited colour, was visited significantly more often than purple and white (p < 0.05, table S5), the least visited colours, but no other pairwise comparisons were significant.

There was no significant effect of colour on visits for greenfinch (F_7,140_ = 1.3.383, p = 0.217) or starling (F_7,84_ = 1.232, P =0.294), and no other species was recorded more than 100 times during the sample period, so their preferences have not been analysed.

**Fig 2:**
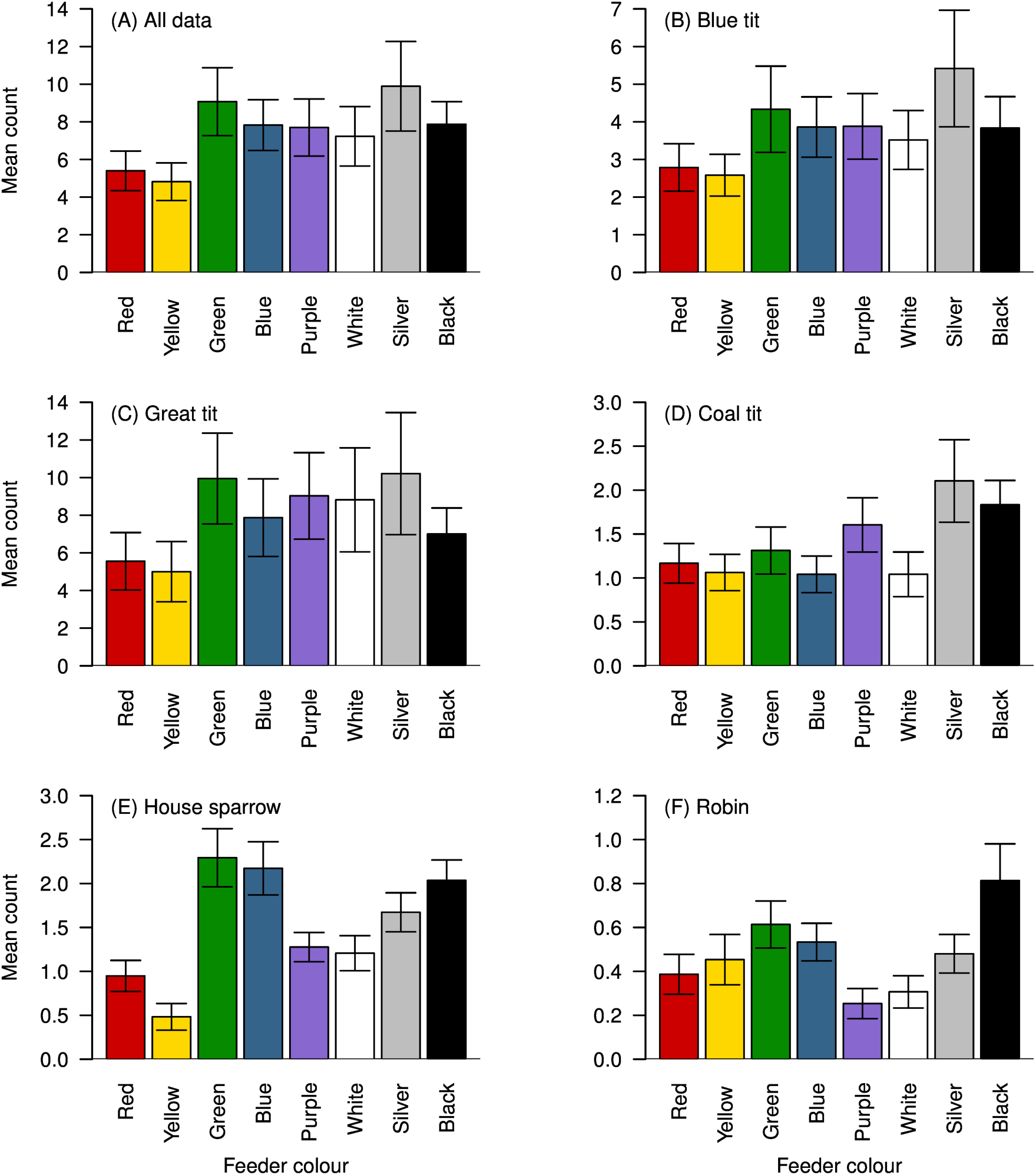
Bird colour preferences. Mean numbers of visits per observation period to feeders ofeach colour, for **(A)** all species combined, **(B)** Blue tit *Cyanistes caeruleus* **(C)** Great tit *Parus major* **(D)** Coal tit *Periparus ater* **(E)** House sparrow *Passer domesticus* and **(F)** Robin *Erithacus rubecula*. Error bars represent +/−1 S.E.

### Human colour preference

There was a significant effect of colour on the preferences observed in our survey (F_1,7_=10.485, P=<0.001; Fig 3a). Pairwise comparisons revealed that red, yellow, green and blue were preferred over purple, white, silver and black (table 3). Fig 3B shows the mean number of visits by birds plotted against the mean number of votes from visitors, and suggests that human and bird preferences do not necessarily align. Colours in the top right of Fig 3B are those that received high visit rates from birds and high numbers of votes from visitors, and we suggest those colours (green and to a lesser extent, blue) may be simultaneously marketable and well-visited by birds. While red and yellow received high numbers of votes from visitors, these are the colours that received the lowest numbers of visits from birds.

**Table 3:** Pairwise comparisons of votes by visitors to the garden centre and science festival.

|  | Red | Yellow | Green | Blue | Purple | White | Silver | Black |
| --- | --- | --- | --- | --- | --- | --- | --- | --- |
| Red | - | z = -0.474<br>p = 0.810 | z = 0.620<br>p = 0.715 | z = -0.182<br>p = 0.0.887 | z = 3.285<br>**p = 0.003** | z = -4.014<br>**p < 0.001** | z = -4.234<br>**p < 0.001** | z = 3.759<br>**p = 0.001** |
| Yellow | -0.722±1.522 | - | z = 0.146<br>p = 0.884 | z = -0.657<br>p = 0.718 | z = 2.810<br>**p = 0.011** | z = 3.540<br>**p = 0.002** | z = 3.759<br>**p = 0.001** | z = 3.285<br>**p = 0.003** |
| Green | 0.944±1.522 | 0.222±1.522 | - | z = -0.803<br>p = 0.659 | z = -2.664<br>**p = 0.015** | z = -3.394<br>**p = 0.002** | z = -3.613<br>**p = 0.001** | z = 3.139<br>**p = 0.004** |
| Blue | -0.278±1.522 | -1.000±1.522 | -1.222±1.522 | - | z = -3.467<br>**p = 0.002** | z = -4.197<br>**p < 0.001** | z = -4.416<br>**p < 0.001** | z = 3.942<br>**p < 0.001** |
| Purple | 5.000±1.522 | 4.278±1.522 | -4.056±1.522 | -5.278±1.522 | - | z = -0.730<br>p = 0.688 | z = -0.949<br>p = 0.568 | z = 0.474<br>p = 0.810 |
| White | -6.111±1.522 | 5.389±1.522 | -5.167±1.522 | -6.389±1.522 | -1.111±1.522 | - | z = 0.219<br>p = 0.891 | z = -0.255<br>p = 0.895 |
| Silver | -6.444±1.522 | 5.722±1.522 | -5.500±1.522 | -6.722±1.522 | 1.444±1.522 | 0.333±1.522 | - | z = -0.474<br>p = 0.810 |
| Black | 5.722±1.522 | 5.000±1.522 | 4.778±1.522 | 6.000±1.522 | 0.722±1.522 | -0.389±1.522 | -0.722±1.522 | - |

The cells above the diagonal show the z- and p-values, while the estimate ± standard error is below the diagonal. Significant p-values are highlighted in bold.

**Fig 3:**
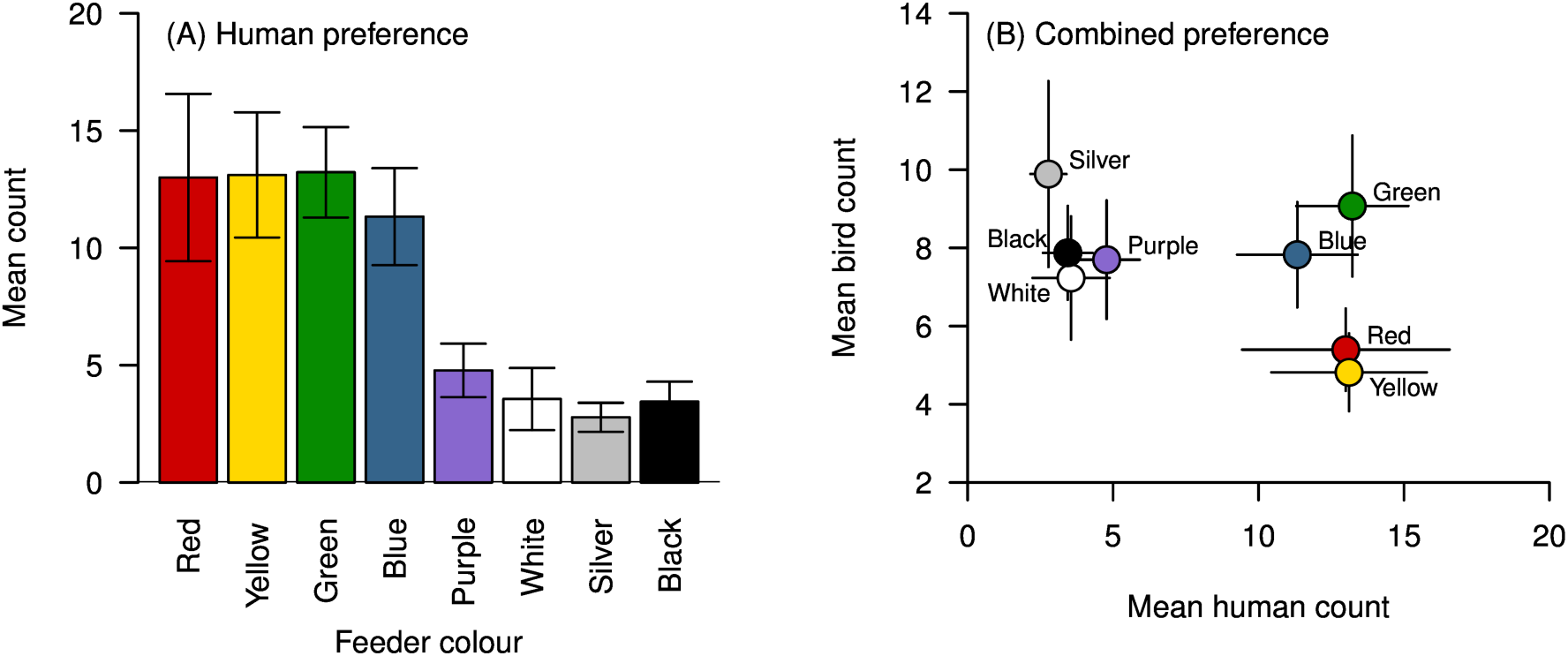
Human colour preferences. **(A)** Mean number of tokens placed into the container corresponding to each coloured feeder by potential purchasers of bird feeders. **(B)** The combined preferences of potential purchasers (x axis) and visits by all birds (y axis) for each colour feeder. Error bars represent +/- 1 S.E.

## Discussion

Overall, birds preferentially visited the silver feeders, followed by green, and made fewer visits to the red and yellow feeders when all feeders were displayed simultaneously. These patterns are likely driven by the preferences of the most abundant species at the feeders (blue tits and great tits), which showed similar preferences to the overall patterns. These patterns contrast with the preferences expressed by the potential purchasers of feeders, who preferred red, yellow, green and blue, but rarely voted for silver. In terms of feeder design, this suggests that green (and to a lesser extent, blue) may be simultaneously marketable and well visited by birds. Our findings also suggest that different species of birds may have different colour preferences, although the total number of visits by some species was too low to evaluate this.

Silver and green feeders may be preferred over red and yellow for a variety of reasons. Green and silver are common colours for birdfeeders, and familiarity with particular colours may have played a role in determining preferences. In hummingbirds, previous experience of particular colours plays a role in colour choice. Anna’s (*Calypte anna*) and rufous (*Salasphorus rufus*) hummingbirds preferentially choose red feeders if captured from red-flowered *Ribes speciousm* plants, but prefer yellow if captured near yellow-flowered *Nicostiana glauca* [38]. Hummingbirds can also be trained to prefer particular colours when that colour is associated with higher quality rewards [31, 35, 39]. As the seed quality in our feeders was identical, the preferences exhibited by the birds could have been due to our choice of locations where birds were regularly fed, and thus familiar with commonly coloured feeders.

In contrast, red and yellow are uncommon colours for seed dispensing bird feeders. Neophobia in relation to food colour has been well documented in both birds (e.g. [47–51]) and other species [52, 53]. Red and yellow are also associated with warning colouration and aposematism, and may be avoided by foraging birds [54,55]. Red and yellow feeders may also be more conspicuous against the background (while green and silver are more cryptic), which may increase perceived predation risk [56]. However, these colours may also make the resource more conspicuous from a distance [57] and thus brightly coloured feeders may be effective in attracting birds to new foraging sites more rapidly: some evidence from Anna’s hummingbirds suggests that red feeders placed in novel locations initially attract more birds than other colours [36], but red is a common colour of the nectar resource for this species, and so may not be applicable to seed-feeding birds. Our experiment does not allow us to speculate on whether particular colours would be more or less attractive to birds if put out alone.

During data collection, we observed (but did not record) multiple events where a competitor displaced feeding individuals from one feeder to another. This may mask feeding preferences as individuals are then recorded at less preferred feeder colours, a limitation of presenting all colours together. The ways in which different options are presented often affects the choices that animals make. Hummingbirds offered a choice between red and yellow-flowered *Mimulus aurantiacus* prefer to feed at the red morph [58], but in a hybrid population where orange morphs occur, visit orange flowers more often than expected by chance, given their prevalence in the population [28]. Preferences between two option may also be affected by the addition of a third option (e.g. if A is preferred over B, and B over C, then A is not necessarily preferred over C), violating the principle that choices are ‘rational’ and preferences are transitive [59,60]. Evidence suggests that the principle of rational choice is violated by a range of species, including humans (e.g. [61–63]), honeybees (*Apis mellifera* [64]), rufous hummingbirds [59,65], starlings [66] and grey jays (*Perisoreus canadensis* [64]). By presenting all colours together (and covering a wide range of colour options) we were able to overcome some of these issues associated with animal decision-making.

Further work is needed to explore interspecific differences in colour preference: for the bird feeder industry, it may be desirable to design and market feeders for particular target species or groups of species - those that are seen as desirable by the people that feed birds. Further questions include whether feeder colour is important for attracting birds to a new feeding site, increasing avian visitor numbers at existing feeding sites, and whether different types of feeders, such as those designed to dispense seeds, peanuts or nyjer seeds, would attract more birds if they were different colours. Finally, other factors, such as distance to cover, or the type and quality of food provided, may be more important in determining the ‘success’ of a particular feeder than the colour, as in hummingbirds (e.g. [35–39]) or these factors, and others, may trade off with colour in determining the number of visits by birds, as they forage optimally [67].

## Acknowledgements

We thank Rose Bull and Adam Lea for assistance with data collection, William Allen for assistance with the feeder colour analysis, Martin McDaid and Lorron Bright for useful discussions, and Hornsea Garden Centre for allowing us to collect data on human choices. Roland Ennos, Sue Hull and Katherine Jones provided useful feedback on earlier versions of this manuscript.

### Supporting information

**Fig S1:**
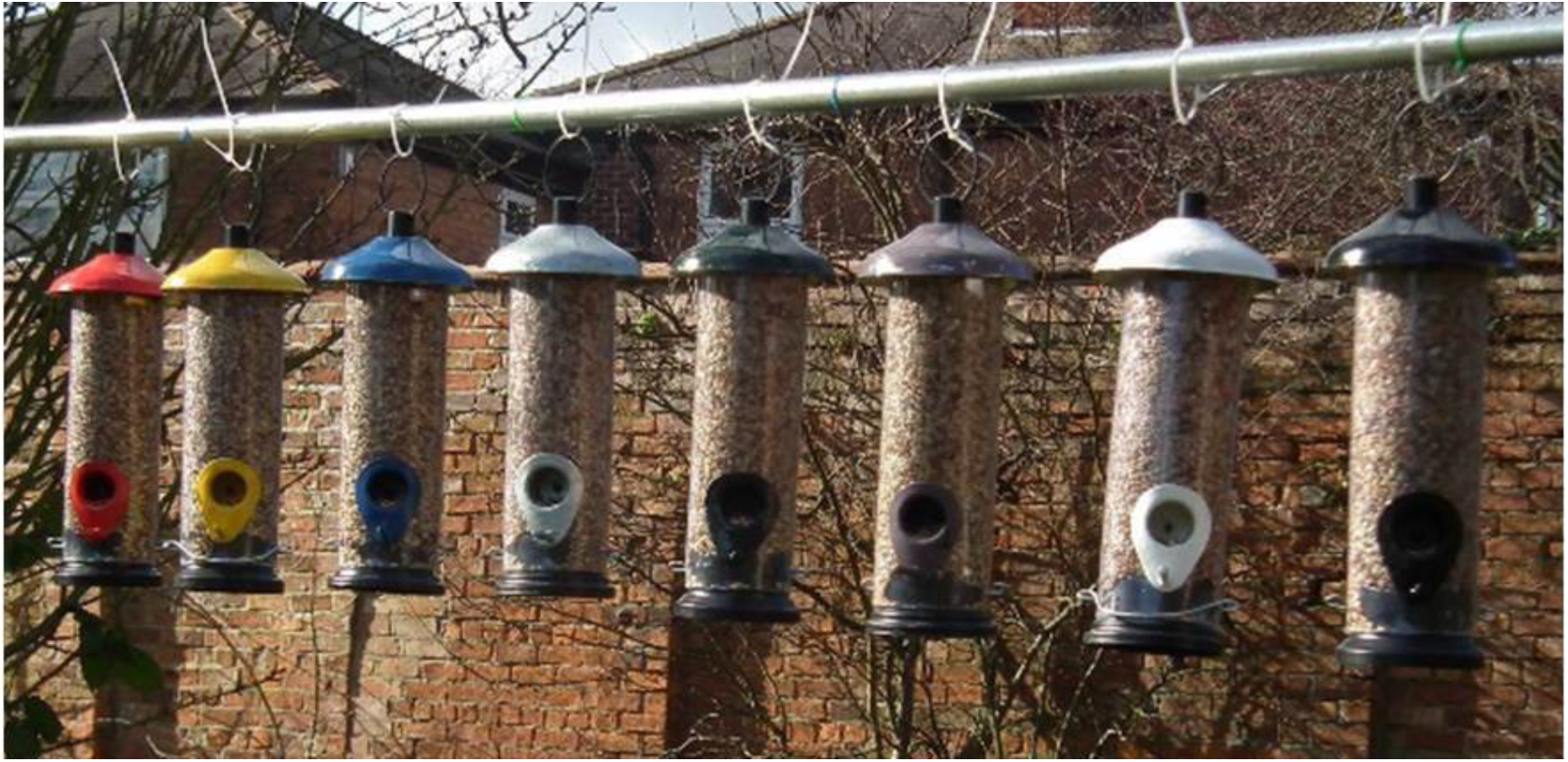
Birdfeeders in the field. An example array of filled birdfeeders ready for observations in the field. The colour order (from left to right) is: red, yellow, blue, silver, green, purple, white, black. Colour order was randomised between trials.

**Fig S2: Poster explaining the project.** A copy of the poster explaining the project, as displayed at the Science Festival and in the garden centre.

**Table S1:** Pairwise comparisons of visits to feeders by blue tits. The cells above the diagonal show the z- and p-values, while the estimate ± standard error is below the diagonal. Significant p-values are highlighted in bold.

**Table S2:** Pairwise comparisons of visits to feeders by great tits. The cells above the diagonal show the z- and p-values, while the estimate ± standard error is below the diagonal. Significant p-values are highlighted in bold.

**Table S3:** Pairwise comparisons of visits to feeders by coal tits. The cells above the diagonal show the z- and p-values, while the estimate ± standard error is below the diagonal. Significant p-values are highlighted in bold.

**Table S4:** Pairwise comparisons of visits to feeders by house sparrows. The cells above the diagonal show the z- and p-values, while the estimate ± standard error is below the diagonal. Significant p-values are highlighted in bold.

**Table S5:** Pairwise comparisons of visits to feeders by robins. The cells above the diagonal show the z- and p-values, while the estimate ± standard error is below the diagonal. Significant p-values are highlighted in bold.

